# The activation of CT fibers or affective touch does not affect pain sensitization by secondary hyperalgesia

**DOI:** 10.64898/2026.01.27.701964

**Authors:** Márcia da-Silva, A. Ribeiro-Carreira, Mariana Oliveira, Adriana Sampaio, Joana Coutinho, Alberto J. González-Villar

## Abstract

Interpersonal affective touch, that preferentially engages C-tactile (CT) afferents, has been shown to produce analgesic effects, yet its role in pain sensitization remains poorly understood. This study explores whether touch delivered at CT-optimal parameters—either artificial or interpersonal—modulates secondary hyperalgesia (SH) induced by high-frequency stimulation (HFS). Two experimental studies were conducted. In Study 1, 46 participants underwent SH induction during two conditions: robot stroking at CT-optimal velocities and vibrotactile stimulation. In Study 2, 64 participants (32 couples) experienced SH in two separate sessions: alone or accompanied by their partner delivering affective touch. Pain reports, electroencephalographic activity (N-P complex and time-frequency activity), electrocardiographic (ECG) and electrodermal activity data (EDA) were collected.

HFS successfully established SH in both studies; however, no significant differences were found in the SH area between CT-stimulation and control conditions. In Study 2, partner-delivered affective touch significantly reduced reported acute pain during HFS compared to the alone condition, whereas no such effect was observed in Study 1 (robotic vs. vibrotactile). At the neural level, no condition effects were observed in the N-P complex, EEG time–frequency data and ECG indexes in either study. Tonic EDA after HFS was higher when participants received stroking from their romantic partner compared to being alone. No sex differences were observed.

Overall, while affective touch from a romantic partner may reduce acute pain during nociceptive stimulation -an effect not observed with robotic CT-stimulation- neither robotic nor interpersonal affective touch appeared to modulate the development of secondary hyperalgesia.

## Introduction

Nociceptive signaling is primarily mediated by unmyelinated C-fibers and typically associated with negative valence. By contrast, gentle touch is often perceived as pleasant and is thought to preferentially engage a specialized subset of C-fibers known as C-tactile (CT) afferents, predominantly found in hairy skin [34,41,58]. According to the “social touch” hypothesis, CT-afferents are thought to contribute to the pleasant subjective experience that occurs alongside autonomic, hormonal, and behavioral responses to gentle touch [46]. Even in the absence of explicit social cues, signals conveyed via CT-receptors are often interpreted as positive, being an efficient method of transmitting embodied (non-verbal) social support [24]. Touch also plays a central role in intimate romantic relationships [55], with perceived pleasantness increasing with relationship quality [57]. Beyond hedonic appraisal, psychological intimacy seems to mediate the regulatory role of touch by communicating a sense of connection, contributing to better affect and well-being over time [12].

Stroking applied prior to a noxious stimulus has been shown to reduce pain, when applied by a romantic partner [42] or by a stranger [25]. Analgesic effects of CT-targeted touch were also observed during simultaneous application with nociceptive stimulation in acute pain paradigms and in protocols modeling central sensitization, such as Temporal Summation Pain [16,33,52]. Although these findings indicate that affective touch can modulate pain under conditions involving heightened nociceptive stimulation, far less is known about whether such social-affective mechanisms extend to a more persistent manifestations of central sensitization. In this context, secondary hyperalgesia (SH) provides a particularly relevant model. SH refers to the heightened pain sensitivity in the uninjured areas surrounding a tissue injury, reflecting increased responsiveness of nociceptive neurons in the central nervous system [4,54]. SH is widely recognized as a robust manifestation of central sensitization that can be induced under controlled experimental conditions [32]. Previous research on SH and social support have shown that handholding can reduce the area of SH, although psychophysiological measures failed to detect this social effect [21].

In this paper, we aimed to investigate how different types of CT-tactile stimulation and social contexts modulate the establishment of SH through two experimental studies. The first study examined the effects of robot brushing compared to vibration, allowing us to isolate the contribution of CT-afferent activation from that of other low-threshold mechanoreceptors (Aβ fibers). The second study explored the effects of interpersonal affective touch in romantic couples compared to the absence of social support. Given evidence for sex differences in touch perception [20,51], but most studies focused exclusively on women [10,17,18,26,35,38,42], in this study we explored sex differences using a balanced sample of men and women.

In both studies we combined a multimodal set of measures: subjective pain reports, SH area, electrophysiological measures of nociceptive processing (pinprick-evoked potentials and EEG time-frequency activity) and autonomic measures of arousal and regulation (electrodermal activity and cardiac indices).

Together, these studies clarify how different tactile interactions and social contexts affect the establishment of SH and the related neural and psychological responses.

## Methods

### Participants, ethics, and preregistration

#### Study 1

A total of 46 participants aged between 18 and 43 years old (*M* = 20.8; *SD* = 5.9; 4 male, 45 female) participated in the study. All participants were students at the University of Minho and received academic credits for their participation. One participant did not provide their date of birth; hence this individual’s data was excluded from descriptives. Additionally, four participants were excluded from all the analysis because during the task they removed the arm from the robot device and did not receive the robot stroking.

#### Study 2

A total of 64 participants (32 couples) aged between 18-41 years old (*M* = 25,3; *SD* = 5,7; 33 male, 31 female) were involved in the study. Participants were recruited through the platform of credits from the University of Minho, social media advertising posters and between participants (snowball approach). University students received course credits, while non-student participants were rewarded with vouchers. Participants must have been in a stable romantic relationship for at least one year.

Exclusion criteria for both studies included: previous history of cardiac, psychiatric, or neurological conditions, history of chronic pain, substance abuse, pain on the day of the experiment, and injured skin on the forearms or hands.

Prior to the experiment, all participants signed an informed consent, acknowledging the voluntary nature of their participation and their right to withdraw at any time. Participants were naïve to the true hypothesis of the study. Participants were informed that they would take part in a study involving moments with painful somatosensory stimuli induced by electrical stimulation, with the aim of examining the neural mechanisms underlying these responses. The study was conducted in accordance with the Declaration of Helsinki and received approval by the Ethics Committee for Social and Human Sciences of the University of Minho (CEICSH 030/2022). Both studies were preregistered on the Open Science Framework before data collection (study 1- https://osf.io/ahnj3; study 2- https://osf.io/dz29k).

### Experimental procedure

When arriving at the laboratory, participants were informed about the study protocol and signed the informed consent. Then, they moved to a room where the electrode cap for electrophysiological recording was placed. During this period, they completed a sociodemographic questionnaire. During the experiment, participants were comfortably seated in a chair in a room with controlled lighting and soundproofing. Prior to the experimental protocol, a period of 4 minutes of resting state with eyes open was recorded.

#### Mechanical Pinprick Stimulation

The experimenter informed the participants that they would receive pinpricks in their forearms and that they needed to remain steady and keep their eyes on the screen. The device was shown before the EEG collection to guarantee that they were familiar with it. The pinprick stimulator was composed of a 128mN stainless-steel flat-tip probe of 0.25 mm diameter that moves freely inside a handheld stainless-steel tube and is capable of sending triggers to collect evoked brain activity (MRC Systems, Heidelberg, Germany). The tube was held perpendicular to the skin, and the experimenter who was holding it moved it up and down, with approximately 1 second of contact with the skin.

Pinprick evoked potentials on both forearms were recorded prior to (T0) and after (T1) the induction of SH. Eighty-two mechanical pinpricks were applied on each arm (41 in T0 and another 41 in T1), with an inter-stimulus interval of 8-11 seconds. To determine the area of stimulation in T0, a circle with a radius of 2,5 cm was delineated on both participants’ inner forearm arms, positioned about 2 cm below the cubital fossa. The determination of the area of secondary hyperalgesia for performing the pinpricks in T1 is described in the following section.

In each pinprick, the stimulus was applied to a different location inside the area. For every 10 pinpricks, the participant rated the intensity of pain induced by the last pinprick on a numerical pain scale with verbal anchors shown on a computer screen.

#### Induction of secondary hyperalgesia using high frequency electrical stimulation

High-frequency electrical stimulation (HFS) consisted of five 1-second trains of electrical pulses at 100Hz applied to the volar forearm, with one HFS train delivered every 10 seconds. Participants provided verbal pain ratings (VPR) after each train to report the perceived pain intensity.

HFS was applied using an EPS-P10 HFS electrode (MRC-Systems). The device includes a cathode composed of ten tungsten pins, each with a stable diameter of 0.25mm and an anode with 410 mm^2^. The stimulator was connected to a Digimiter DS7A controlled by a DG2A Train Delay Generator. The EPS-P10 was positioned with the cathode placed in the center of the circle. The participant’s detection threshold was determined through the staircase method, starting with a low intensity (.06mV) and gradually increasing it (.1mV step size) until the participant was able to detect the stimulus. After the detection, the intensity was decreased to guarantee the detection threshold. The intensity of HFS was set at 20x the participant’s detection threshold.

The participants were informed about the study conditions and the stimulation they would receive during the high-frequency stimulation. During each stimulation, they verbally reported their pain levels based on the pain scale that was above the computer screen. The scale ranged from 0 to 100, where 0 represented “no sensation”; 10 - “noticeable sensation”; 20 - “mildly painful sensation” (i.e., pain threshold); 30 - “very weak pain”; 40 - “weak pain”; 50 - “moderate pain”; 60 - “slightly strong pain”; 70 - “strong pain”; 80 - “very strong pain”; 90 - “nearly intolerable pain”; and 100 “unbearable pain”. After that, they were instructed to focus on a fixation point shown in the screen for 10 minutes and report their pain levels five more times. The presentation of the HFS trains, pain scales and fixation point was controlled using the software PsychoPy [47].

The area of SH was subsequently measured with a 128mN punctate calibrated probe, by applying stimulation along the four linear trajectories arranged both vertically (proximal to distal) and horizontally (from lateral to medial) in approximately 1cm increments across the stimulated forearm. Stimulation started at the wrist and at the cubital fossa and then at the inner and outer edges of the forearm progressing toward the center of the stimulation site until the participant signaled a noticeable alteration in the pinprick intensity. This point was marked with a makeup pencil, and the measure was collected using a digital caliper. The area of hyperalgesia was calculated using the formula for the area of a diamond in both conditions for the 2 studies. Finally, the T1 pinprick evoked potentials were recorded again over the area of SH.

#### Study 1

In this study, participants were exposed to a secondary hyperalgesia (SH) protocol in two different conditions: SH during robot stroking and SH during vibrotactile stimulation. The order of presentation of these conditions was counterbalanced among the participants. The arm receiving the initial pinpricks was the first to be stimulated (see fig.1).

**Fig. 1.**
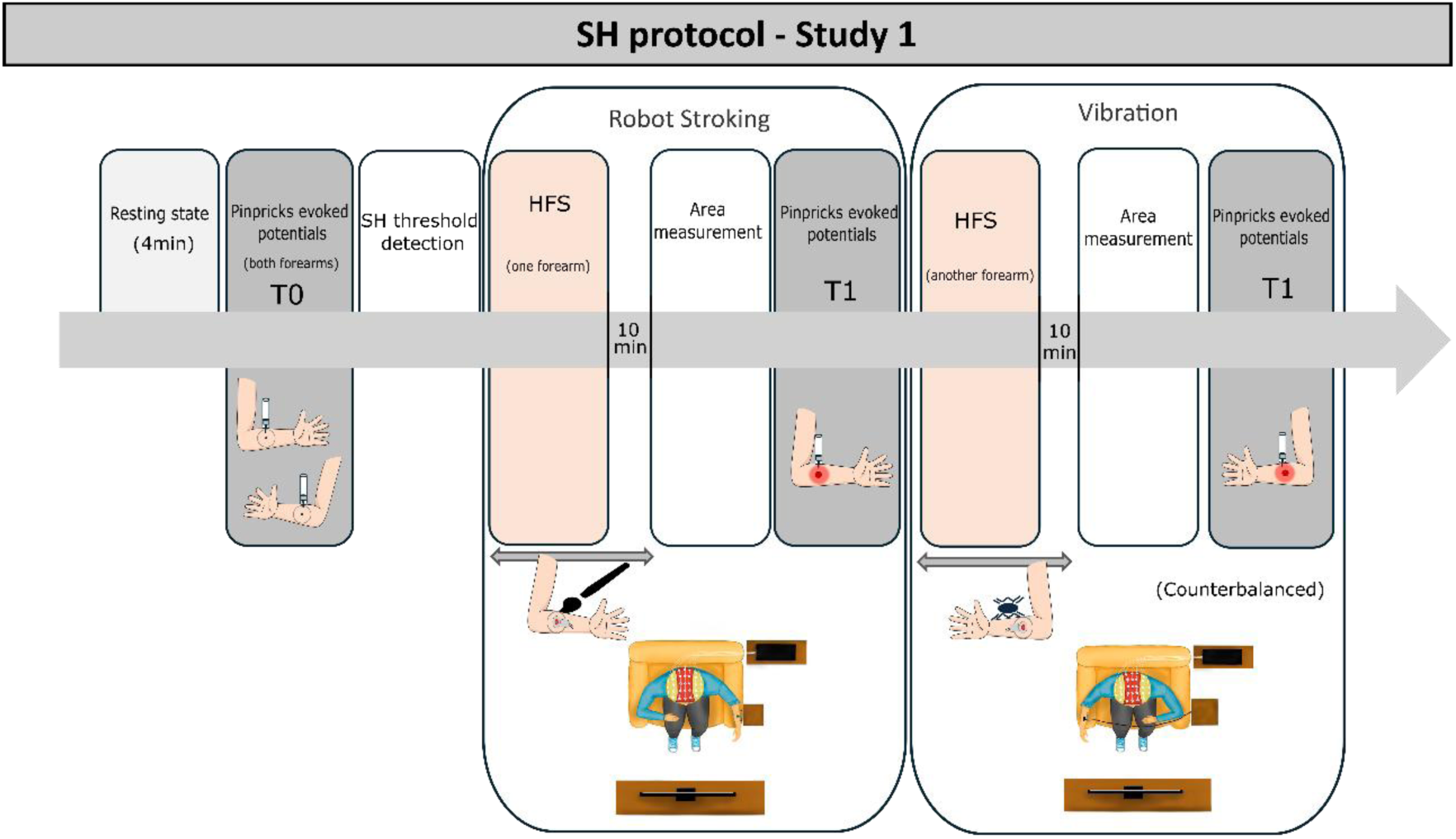
Schematic representation of study 1. Participants underwent a secondary hyperalgesia (SH) protocol using High Frequency Stimulation (HFS) in two counterbalanced conditions: robot stroking in the left dorsal forearm and vibrotactile stimulation in the right dorsal forearm during HFS and in the 10 minutes after.

In the robot stroking condition, the participants’ left dorsal forearm was gently brushed during HFS. The brushing was performed within a designated area measuring 14 cm from proximal to distal and 2.5 cm in width, at a velocity optimized for activating C-tactile fibers (≈ 4 cm/s). This stroking was administered using a cosmetic brush attached to the end of a robotic arm. To prevent visual feedback, the robotic apparatus was enclosed in a box, making the movement of the arm invisible to participants. In the vibrotactile condition, the vibrotactile stimulation was applied to the right dorsal forearm at a frequency of approximately 200 Hz, using a linear resonant actuator. The vibrotactile stimulator comprised two small Linear Resonant Actuators housed in a 3D-printed plate measuring 6*2.5 cm, which was secured to the participant’s forearm with adhesive tape. Both the robot stroking and the vibrotactile stimulation were performed during and 10 minutes after HFS. During HFS, vibration or the robot stroking had a duration of 4.5 seconds after 0.5 seconds of each electrical pulse. In the following 10 min, vibrotactile or stroking stimulation was delivered, back and forth, for 4.5 s with intervals of 10 s.

To ensure perceptual equivalence between the two types of tactile input, the intensity of the vibrotactile stimulation was calibrated for each participant before the task. Participants were informed that the intensity of the robot stroking remained constant and that they could only manipulate the intensity of the vibration. A visual analog scale (0 = “minimum” to 10 = “maximum”) was displayed on the screen, and using the computer mouse, they could modify the vibration intensity until they felt it matched the robot stroking intensity, at which point they clicked the “Accept” button. This calibration process was repeated 3 times, and the average intensity of vibration from these 3 trials was set as the task intensity. Participants were allowed to restart and repeat the calibration procedure if they felt they had not calibrated properly.

#### Study 2

The second study was divided into two conditions (dyadic and alone, counterbalanced) that were conducted in separate sessions, with at least one week apart to reduce potential emotional carryover effects.

In the dyadic session, participants were accompanied by their partners throughout the procedure. In this condition, the participant was gently stroked by his/her romantic partner in a pre-defined area of the participant’s arm (14 cm in length) during the induction of SH and for ten minutes afterwards. The stroking area was chosen because it is the same area that was stroked by the robot in Study 1. To ensure consistency across participants and also to guarantee that the velocity matched the desired speed to activate CT-receptors (velocity around 3-5cm/s [1], the speed of the stroking was controlled via auditory cues delivered through headphones to the partner performing the touch. During HFS, each stroking lasted 5 seconds with intervals of 5 seconds of no touch, starting 1 second after the first electrical pulse and ending 6 seconds after final pulse. During the 10 min subsequent HFS the touch lasted 30 seconds (with auditory cues every 3.5 seconds to control for the velocity) separated by 20 seconds of no touch intervals. Before the task, participants’ romantic partners were trained on how to administer the touch, and the researcher provided instructions regarding any deviations from the intended stimuli. The partner only left the EEG room to the adjacent room during the moments of the pinprick stimulation, where the stimulation was performed by the researcher.

In the individual session, participants remained alone during the nociceptive electrical stimulation and for the ten minutes afterwards.

The protocol to induce SH in this study was the same described in Study 1, with the exception that the HFS was applied exclusively in the right forearm.

**Fig. 2.**
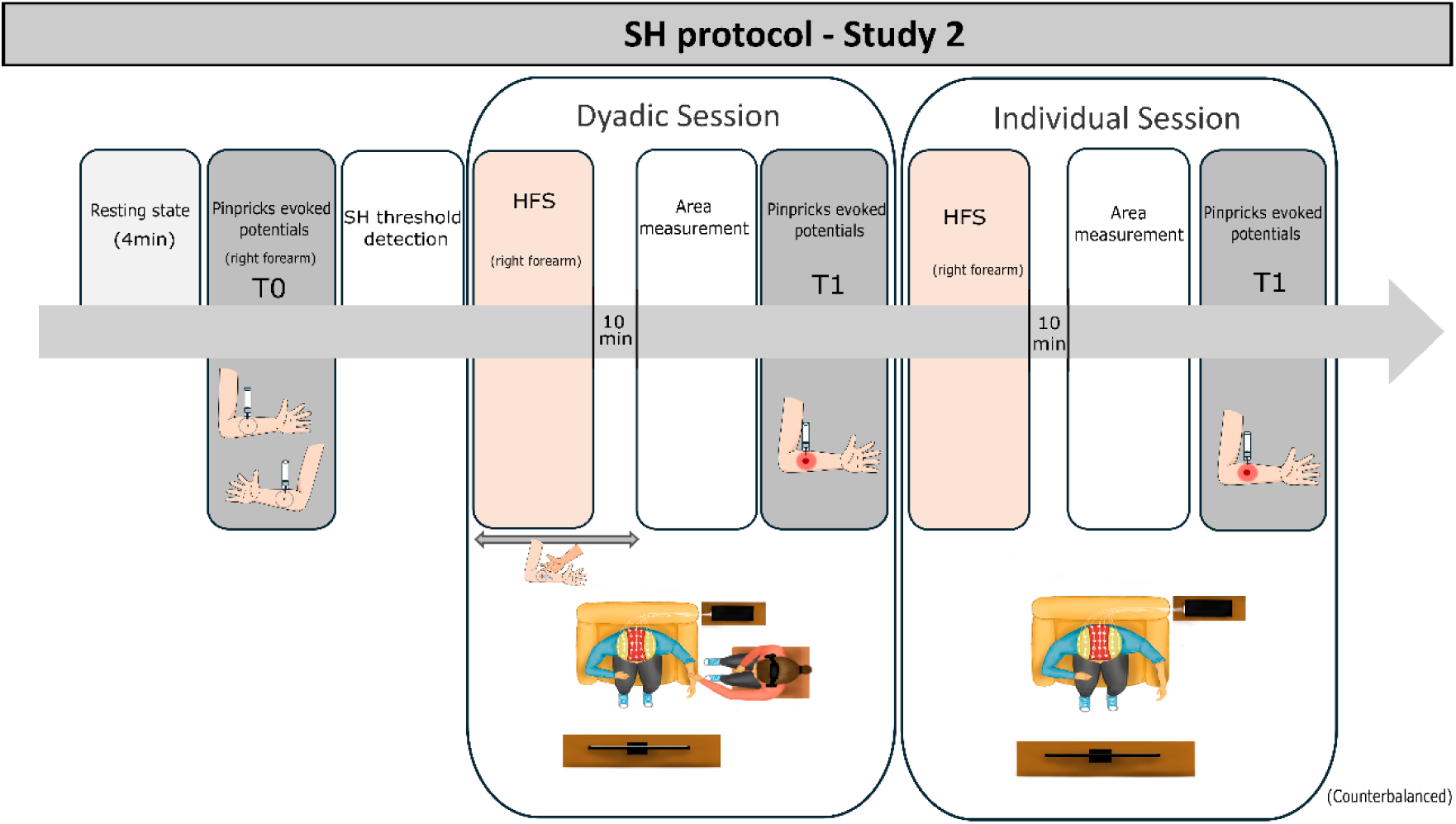
Schematic representation of study 2. Participants underwent a secondary hyperalgesia (SH) protocol using High Frequency Stimulation (HFS) in two counterbalanced sessions: individual and dyadic. In the individual session, HFS was applied to the right volar forearm without additional stimulation. In the dyadic session, participants received HFS on the right volar forearm while their partner provided affective touch on the right dorsal forearm during and for 10 minutes after stimulation.

### Psychological measures

In both studies, participants provided socio-demographic information and filled out the Edinburgh Handedness Inventory to assess their laterality.

They also completed the PHQ-9 questionnaire for screening the presence of depressive symptoms [29,30].

**Fig. 3.**
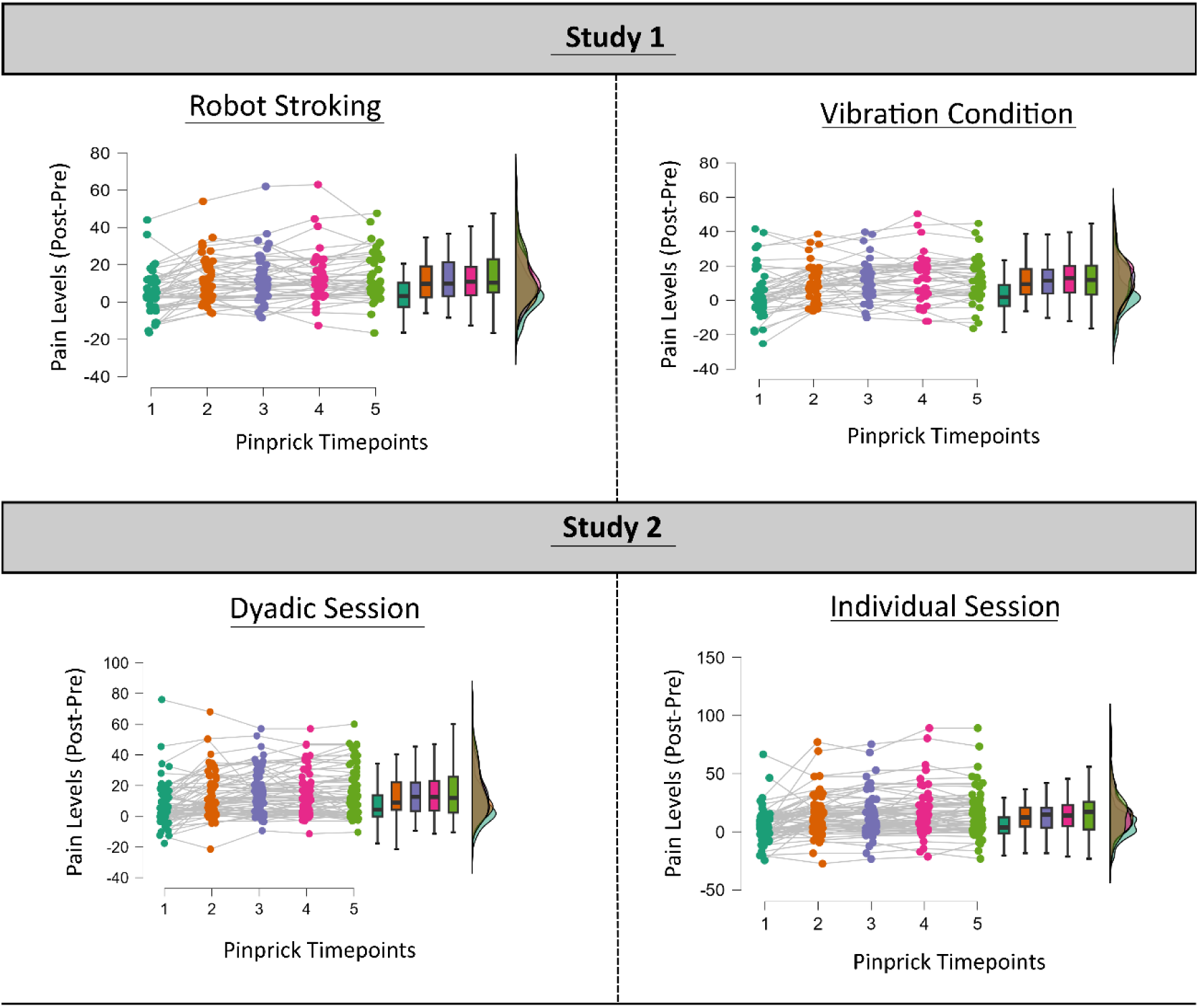
Scatter plot showing pinprick pain levels across both studies under the four experimental conditions. The Y-axis represents the difference in pain after versus before high-frequency stimulation. The X-axis shows five evaluation points: after the first pinprick and after each subsequent set of 10 pinpricks.

### Electrophysiological data

#### Electrophysiological recordings

EEG recording was conducted using the BioSemi ActiveTwo system (Biosemi, Amsterdam, The Netherlands). To capture neural activity, participants wore a nylon cap with 64 active Ag/AgCl scalp electrodes. Electrode locations followed the international standard 10-10 system. Additionally, five active external electrodes were placed in the lateral canthi of both eyes (horizontal electrooculogram - HEOG), below the left eye (vertical electrooculogram - VEOG), and at the tip of the nose. Electrodes were referenced to the Common Mode Sense (CMS) and Driven Right Leg (DLR) Biosemi electrodes. The electrocardiographic activity was recorded by placing an external electrode below the left clavicle. The electrophysiological signals were sampled at a rate of 512 Hz, and an on-line bandpass filter ranging from 0.01 Hz to 100 Hz was applied. Prior to the recording, it was verified if electrode impedances were below 30mV. In both studies, ECG signals were obtained by positioning an electrode below the left clavicle.

In the study 1, Electrodermal activity (EDA) was recorded at 10 Hz using the James two wireless device (MindProber Labs) [53], with electrodes placed on the plantar surface of the left foot. This placement was necessary because participants held the brushing box with their left hand and used the mouse with their right hand to provide subjective ratings; placing the electrodes on the foot helped avoid movement-related artifacts in the EDA data. In study 2, EDA was recorded at 10 Hz using James four wireless device (MindProber Labs) and the electrodes were placed on the palm of the left hand.

#### EEG analysis

The EEG data was analyzed using the EEGLAB toolbox for Matlab (R2022b). An offline bandpass filter was applied to the signals, ranging from 0.1Hz to 30Hz. The data were referenced to the tip of the nose to reduce common noise sources. Epochs of interest were extracted from the continuous data, with a time window of -1000 ms to 2000 ms relative to the onset of the stimulus. Baseline correction from -100 ms to -10 ms relative to the onset was applied. Ocular artifacts were corrected using Independent Component Analysis (ICAs). The data was re-referenced to the average reference. Despite previous studies using a 0.3–30 Hz zero-phase Butterworth filter [7,59,60], we found that a 2-30 Hz band-pass filter improved peak visibility. Before computing the data, we applied a 2-30Hz filter. Results obtained using the 0.3–30 Hz filter are presented in the Supplementary Material.

For pinprick analysis, we computed the average waveform from the 2 conditions in the T0 and T1 moments. For each waveform, the peak amplitudes of N and P components were measured at the Cz electrode. The N component was defined as the most negative deflection following the stimulus onset in a time window from 20 to 100 ms. The P component was defined as the most positive deflection following the stimulus onset in a time window from 100 to 300 ms.

For the time–frequency analysis, we epoched the EEG data in the interval from –990 to 1990 milliseconds relative to stimulus onset. Time-frequency analysis was performed using convolution with complex Morlet wavelets, with frequencies ranging logarithmically from 2 to 30 Hz, spaced into 29 steps. The wavelet convolution was performed in the frequency domain using Fast Fourier Transforms (FFTs) of both the EEG signal and the wavelets, with zero-padding to optimize spectral resolution. The full convolution was computed per trial and electrode, and spectral power was extracted as the squared magnitude of the analytic signal.

Baseline correction was applied by computing the mean power within the –400 to –200 ms pre-stimulus window and converting post-stimulus power values into decibel (dB) changes relative to this baseline. Power was then averaged across trials for each condition. Power data were extracted from electrode Cz for all participants and all conditions. The analysis focused on a broad time window from 0 to 1500 ms and a frequency range of 3 to 30 Hz.

#### ECG analysis

For both studies, ECG analysis was performed using the plugin EEGBeats of EEGLAB in Matlab [56]. The raw EEG data, which included the ECG channel, were imported and preprocessed to remove the stimulation period associated with HFS and the recording up to 2 seconds after this event, ensuring that only post-stimulation resting periods were analyzed. Subsequently, ECG signals were extracted from the continuous data, filtered from 2 to 20Hz, and the R peaks were detected. EEGBeats uses a divide-and-conquer peak detection algorithm that identifies R-peaks directly from raw ECG signals in EEG recordings. It segments the signal multiple times to isolate candidate peaks and applies thresholds based on robust standard deviation, making it robust to noise and artifacts without relying on QRS complex morphology.

For each condition, we obtained the average heart rate (HR), interbeat interval (IBI), and the root mean square of successive differences (rMSSD) of the interbeat intervals. The rMSSD was selected as an indicator of heart rate variability (HRV). These metrics were chosen because HR reflects the average number of heartbeats per minute and is commonly used as an index of physiological arousal, while HRV measures the variation in time intervals between consecutive heartbeats, offering valuable information about autonomic nervous system regulation of cardiac activity [39].

#### EDA analysis

EDA signals were preprocessed and analyzed using custom scripts in Python with the NeuroKit2 library [37]. Raw EDA data were segmented per participant, condition and session (study 2) based on behavioral markers generated during the task. Each segment was decomposed using *NeuroKit’s eda_process function* and from the tonic component we extracted the mean skin conductance level (SCL) during the 10 minutes following the HFS, as an index of sustained autonomic arousal. The focus on the tonic component was chosen because sustained increases in tonic EDA have been shown to reflect prolonged sympathetic activation [8]. Additionally, the tonic component is usually measured over intervals ranging from tens of seconds to hours, while the phasic EDA component is usually measured within 1–6 s after a discrete event [6]. Segments with signal quality issues, such as excessive sweating leading to signal saturation or signal loss, were excluded from further analysis.

#### Statistical analysis

The statistical analysis was performed in the JASP software (version 0.19.0) [22]. A Shapiro-Wilk test was applied to verify the normality of the variables. If the data did not follow parametric distribution we performed alternative tests for non-parametric distribution, such as Wilcoxon signed-rank test

To compare the area of SH between the two conditions, different statistical analyses were conducted in each study. In Study 1, a Wilcoxon signed-rank test was used, as the data did not meet the assumption of normality. In Study 2, a repeated measures ANOVA was performed, with the area of SH across the two conditions as a within-subject factor, and sex as a between-subject factor.

To assess whether SH was successfully induced, we compared the mean reported pain of pinprick stimulation before and after SH induction for both conditions in both studies by performing a Wilcoxon signed-rank test.

To explore the increase in pain levels evoked by the pinpricks, we first computed the mean of the five pre-HFS (T0) pain ratings. Then, for each one of the 5 pinpricks after the HFS (T1), we calculated the difference in pain between the post-stimulation value and baseline values. Since in study 1, the vibration was performed in the right arm and the robot stroking in the left arm; therefore, computing the change from each participant’s own baseline, minimized potential effects of arm laterality, ensuring that the observed differences reflected the experimental manipulations rather than systematic lateralization. In study 2, this subtraction helped us remove any potential effect due to data being collected on different days. To analyze this data, we conducted 2x5 repeated measures ANOVA, where the two conditions (study 1-robot stroking or vibration; study 2- dyadic and individual) were entered as a factor and the five pain ratings (T1-T0) during the pinpricks were entered as another factor. The sex of the participants was entered as between subject factors in study 2. Pinpricks began approximately 12 minutes after the induction of secondary hyperalgesia, whereas the pain levels were reported after the first pinprick and then after every subsequent set of 10 pinpricks. The time between each pain rating varied by approximately one and a half minutes. By entering the five moments of reported pain it is possible to investigate the increase of pain over the period of time that is crucial for the establishment of secondary hyperalgesia. To further explore specific differences between conditions, post-hoc comparisons were conducted using the Holm correction to control multiple comparisons. When the assumption of sphericity was violated, we performed Greenhouse-Geisser correction.

In order to assess differences in pain levels during HFS, we compared the mean values of reported pain across the two conditions after each train of stimulation. In study 1 we performed Wilcoxon signed-rank test and in study 2 a rmANOVA with sex as a between subject factor.

For electrophysiological activity, we computed the amplitude of the peak-to-peak N-P component, and analyzed the difference between before and after the induction of SH (Post minus Pre). For N-P complex and ECG measures we performed a paired sample T-test (Student or Wilcoxon signed-rank) in study 1 and a repeated measures ANOVA in study 2, where the two conditions entered as a factor and the sex as a between subject factors. This enabled the analysis of within-subject effects across the task’s conditions.

For time-frequency analysis, difference maps were then computed for each condition. In the study 1, post-HFS minus pre-HFS power were computed for robot stroking and for vibration conditions; while in study 2, post-HFS minus pre-HFS power were computed for accompanied and for individual conditions. At each time–frequency point, a paired-sample t-test was performed between the two differential conditions across participants, resulting in a map of t-values. Non-parametric cluster-based permutation test was applied to compare between conditions. A total of 10,000 permutations were performed, with a cluster-forming threshold of α = 0.05. Clusters were identified as contiguous time–frequency regions exceeding the significance threshold, and their p-values were computed against the null distribution of cluster-level statistics. Only clusters with corrected p-values below 0.05 were considered statistically significant and visualized on the resulting t-maps. This approach allowed for the identification of temporally and spectrally specific interaction effects without inflating the Type I error rate.

Lastly, as reported on the preregistered analysis plan, we performed Bayesian analysis to improve the reliability and interpretability of the data. Bayes factors (BF_10_) were computed to quantify the relative evidence for the alternative vs null hypothesis. For Bayesian rm-ANOVAs, Cauchy priors were applied to effect sizes (scale = 0.5 for fixed effects, 1.0 for random effects) [43,50]. Bayesian paired-samples t-tests were performed using a Cauchy prior on the standardized effect size (r = 0.707) [50]. The Bayesian analyses were also conducted in JASP.

## Results

### Area of secondary hyperalgesia

In study 1, no differences between conditions (robot stroking or vibration) were observed in the establishment of the SH (z=-0.781, p= .44, Effect Size = -0.138, BF10= 0.284, W= 389, Rhat=1.001, 95% CI= [-0.451,0.141]).

Similarly, the results from study 2 did not show a significant condition effect (F(1,62)=0.028, p=.87,np2= 0.0004, BF10=0.189) nor an interaction effect with sex (F(1,62)=0.078, p=.78, np2= 0.001, BF10=0.097). The area of SH was not influenced by social support or sex of the participant. (See table 1 for the descriptive results).

**Table 1.** Descriptive statistics of SH area for both studies.

| Secondary hyperalgesia area |  |  |
| --- | --- | --- |
|  | Mean (cm<sup>2</sup>) | SD |
| **Study 1** |  |  |
| Robot Stroking | 37.6 | 22.3 |
| Vibration | 40.8 | 28.9 |
| **Study 2** |  |  |
| Dyadic Setting | F=23.5/ M=27.4 | F=11.1/M=12.6 |
| Alone Setting | F=24/ M=26.7 | F=9.1/M=15 |

Note:SD-Standard Deviation; F-Female; M-Male

#### Pre-SH vs Post-SH Pain Comparison

In both studies, mean pain levels reported during pinprick test were higher after HFS than before, across all conditions (see Table 2), indicating that SH was successfully induced in both studies.

**Table 2.**
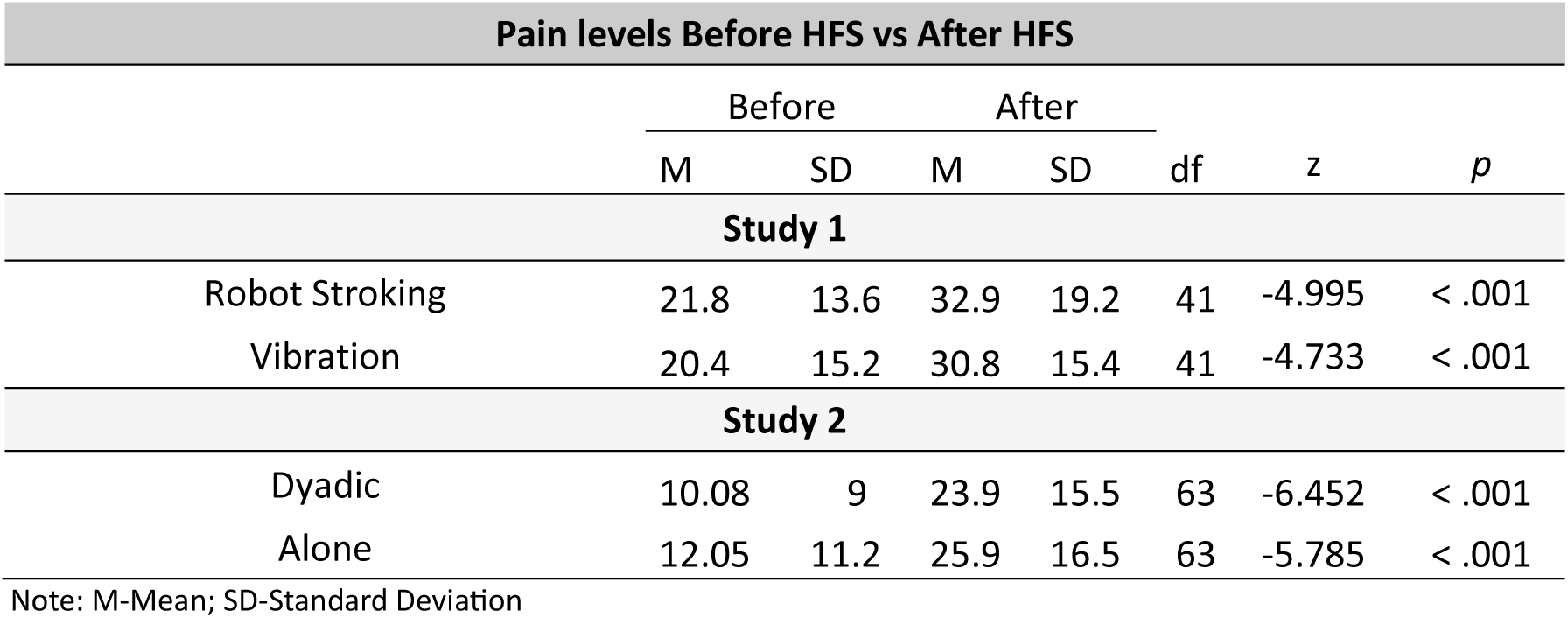
Results of the Wilcoxon signed-rank test for both studies.

#### Acute Pain During HFS

In study 1, no pain differences were observed during HFS stimulation when it was applied while participants were being stroked or compared to when they were receiving vibration (Z=0.45, *p*=0.66, Effect Size = 0.082, BF_10_=0.180, W=422.0, Rhat=1.00, 95%CI= [-0.231,0.352]). In study 2, receiving stroking from a romantic partner reduced reported acute pain during high-frequency stimulation compared to being alone (F(1,62)=7.039, *p*=.010, n_p_ ^2^=0.1, *p*_holm_ =.010, Cohen’s d= -0.3, BF_10_=4.1), with no interaction with sex (F(1,62)=0.011, *p*=.9, n_p_ ^2^=0.0002, BF_10_=2). (See table 3 for the descriptive results).

**Table 3.** Descriptive statistics of acute pain levels during HFS for both studies.

| Acute pain levels |  |  |
| --- | --- | --- |
|  | Mean | SD |
| Study 1 |  |  |
| Robot Stroking | 78.3 | 15.2 |
| Vibration | 77.4 | 16 |
| Study 2 |  |  |
| Dyadic Setting | $F=72.2/ M=68.2$ | $F=19.7/M=16.5$ |
| Alone Setting | $F=77/ M=73.4$ | $F=16,3/M=15$ |

Note:SD-Standard Deviation; F-Female; M-Male

Lastly, we did not observe any differences in the intensity of the current used to perform the HFS between the two sessions in study 2 (F(1,62)=2.66, *p*=.1, n_p_ ^2^=0.041, BF_10_=0.61), but we observed an interaction between the value of the HFS intensity and sex (F(1,62)=4.33, *p*=.04, n_p_ ^2^=0.065, BF_10_=3.5). Post hoc comparisons indicated that male participants had significantly higher HFS values when alone compared to when accompanied (*p* = .039). A significant difference was also observed between females and males in the individual condition (*p* = .028). Lastly, even in the dyadic conditions, females had a lower HFS values compared to male participants in the individual condition (*p*=.031). No other comparisons reached statistical significance after correction.

#### Pain Increase During Pinprick Stimulation

In Study 1, no effect of condition was observed (F(1,41)=0.159, *p*=.69, n_p_^2^=0.004, BF_10_=0.48). Pinprick-evoked pain levels did not differ between the robot stroking and vibration conditions. However, there was a significant effect of time following the induction of SH (F(2.2,91.7)=20.05, *p<*.001, n_p_^2^=0.33, BF_10_=8.2x10^10^). Post Hoc tests indicated that pain for the first pinprick was significantly lower than for the subsequent four pinpricks, with no significant differences among these four (*p*<.001 for all comparisons).

In study 2, no main effect condition was found (F(1,62)=3.9x10^-4^, p=0.98, n_p_^2^=6.3x10^-6^ BF_10_=0.239) nor an interaction with sex (F(1,62)=0.031, *p*=.86, n_p_^2^=0.0005, BF_10_=0.05), suggesting that sex did not influence pain levels during pinpricks. However, there was a significant effect of time (F (1.77, 109.9)=32.8, *p<*.001, n_p_^2^=0.35, BF_10_=3.9x10^18^). Similar to study 1, post hoc tests showed that pain during the first pinprick was significantly lower than during the subsequent four, with no differences among these four (*p*<0.001 for all comparisons).

### Electrophysiological data

#### N-P complex

Due to excessive movement, data from two participants in study 1 (both female) and from three participants in study 2 (all male) were excluded from the analysis. Consequently, electrophysiological data were analyzed from 40 participants in Study 1 and 61 participants in Study 2.

In study 1, no differences in the N-P complex (calculated as the subtraction between pre-and post-induction of SH) were observed between the robot stroking and vibration conditions (z=-1.64, *p*=.1, Effect Size=-0.3, BF_10_=1, W=288, Rhat=1).

Similarly, ANOVA results from study 2 revealed no significant differences in the N-P complex amplitude (also calculated as the subtraction between pre- and post-induction of SH) between individual and dyadic settings (F(1,59)=0.063, *p*=.8, n_p_^2^=0.001, BF_10_=0.198). A significant interaction between condition and sex was observed (F(1,59)=5.016, *p*=.029, n_p_^2^=0.078, BF_10_=0.084), however, post hoc tests did not reveal any significant differences between sexes within either condition (see table 4). Additionally, results demonstrated weak evidence in favor of the null hypothesis for sex differences (BF_10_=0.428).

**Table 4.**
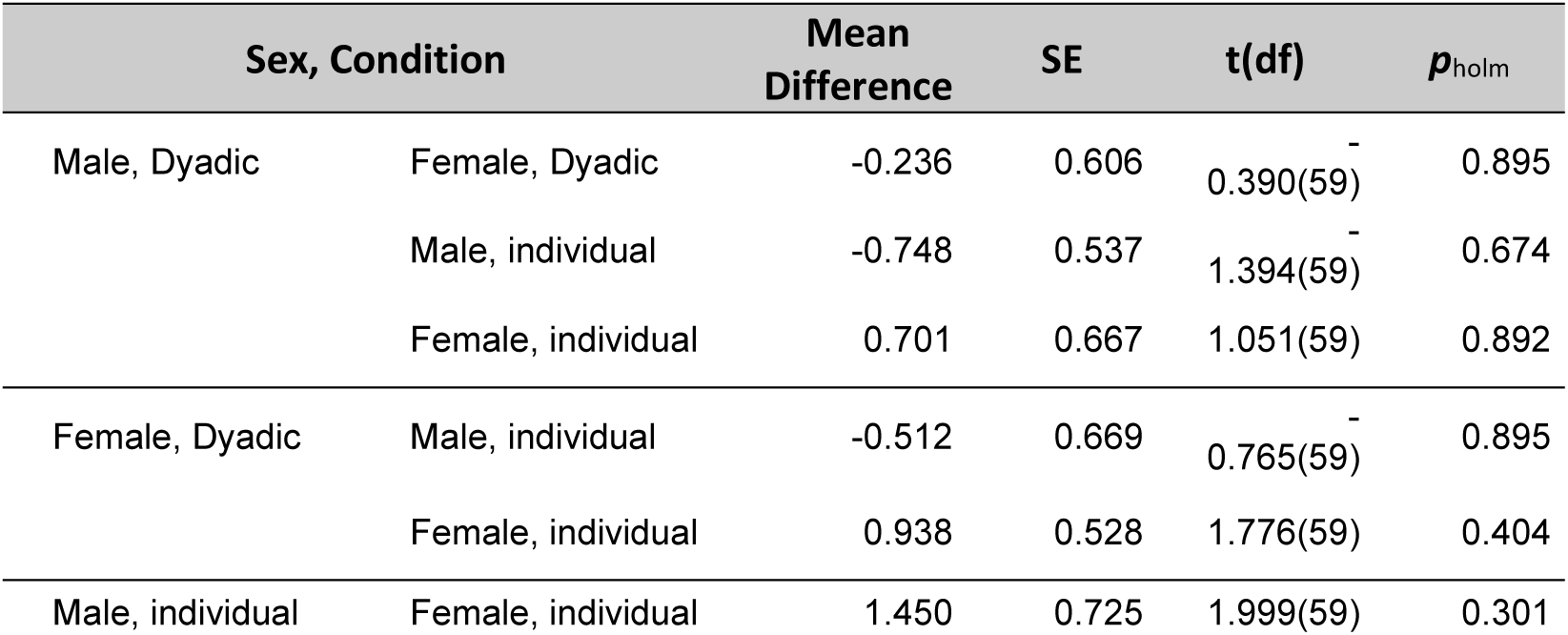
Results from Post Hoc tests of the interaction between condition and sex in study 2.

**Fig 4.**
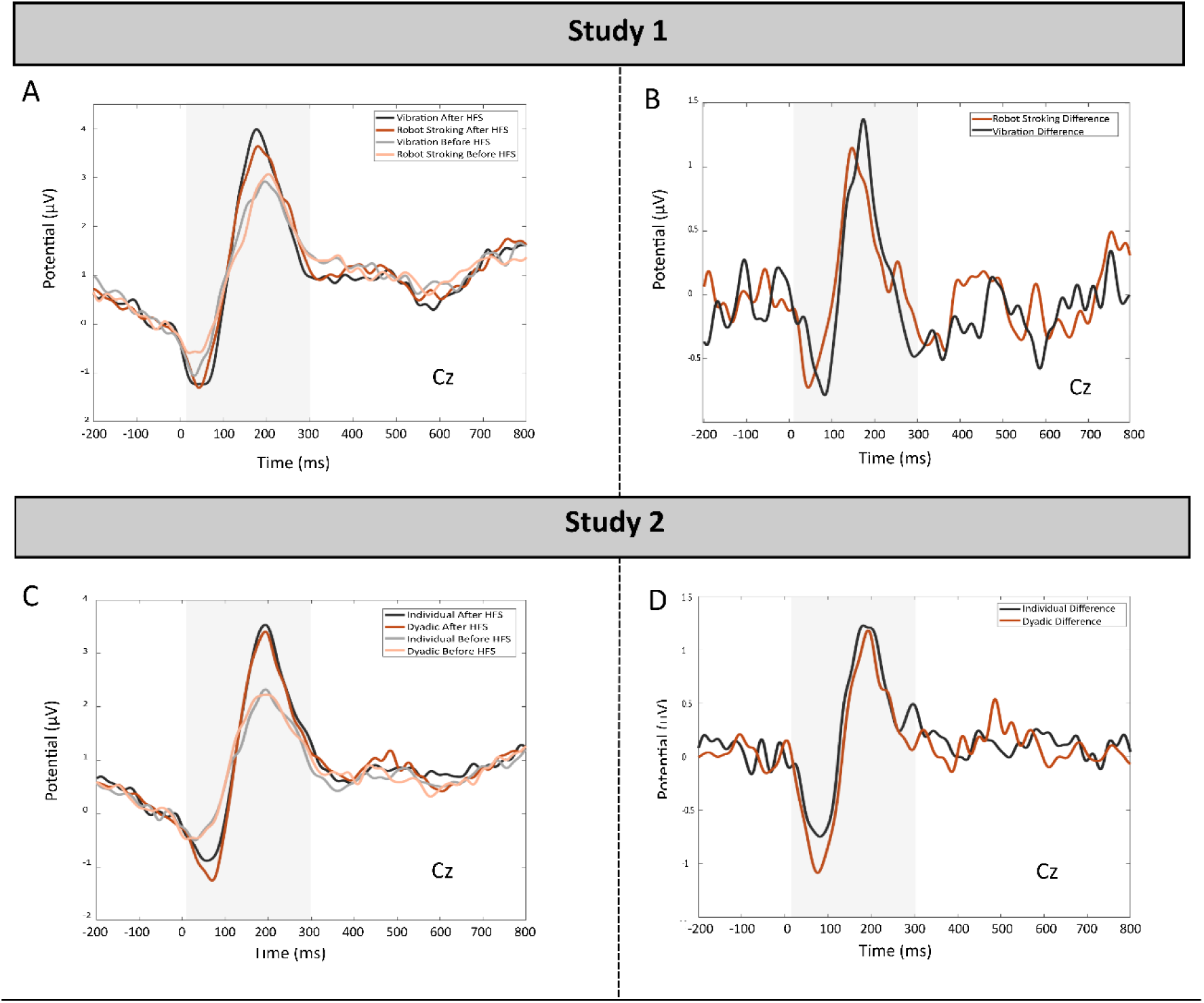
Pinprick-evoked potentials recorded over the CZ electrode in both studies. **Study 1** (A, B): A-Pinprick-evoked potentials at baseline (T0) and post-induction (T1) for Robot stroking and Vibration conditions; B-Difference waveform (T1 minus T0). **Study 2** (C, D): C-Pinprick evoked potentials at T0 and T1 for Dyadic and Individual conditions; B- Difference waveform. HFS- High Frequency Stimulation.

#### Time frequency data

As observed in fig.5, results from time frequency analysis did not reveal any significant clusters from the permutation analysis in both studies.

**Fig.5.**
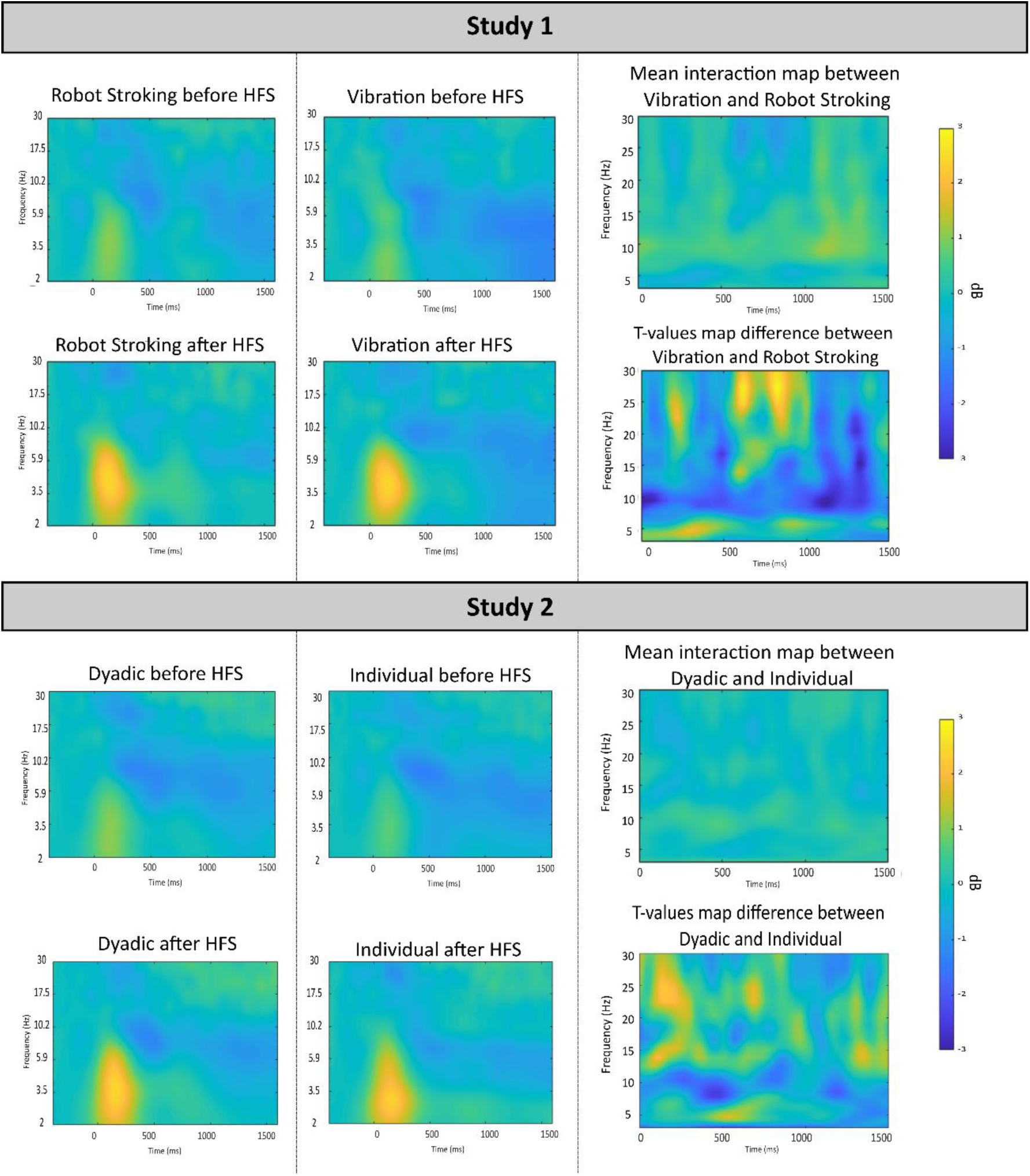
Spectrograms obtained from time-frequency analysis of the pinprick evoked potential, measured at the Cz electrode. Top (Study 1): (Left) Time-frequency plots for each condition before and after HFS. (Right) Mean time–frequency interaction map comparing Vibration and Robot Stroking conditions with corresponding T-values map showing their differences. Bottom (Study 2): (Left) Time-frequency plots for each condition before and after HFS. (Right) Mean time–frequency interaction map comparing Dyadic and Individual conditions, with corresponding T-values map showing their differences. Color scales represent power values (interaction maps) and T-values (statistical maps) across time (0–1500 ms) and frequency (1–30 Hz).

#### ECG

In Study 1, t-test results revealed no significant differences between conditions in any of the ECG measures (Heart Rate, Root Mean Square of Successive Differences and Ratio of Low-Frequency to High-Frequency power of Heart Rate Variability) during the 10 minutes following SH induction. Similarly, Study 2 results showed no significant differences in any of these ECG measures during the same time period, nor were any interactions with sex observed (see Table 5).

**Table 5.** Results of electrocardiographic indexes from studies 1 and 2.

| Study 1 |  |  |  |  |  |
| --- | --- | --- | --- | --- | --- |
| ECG measures | t(sd)/W | Z | p | Effect Size |  |
| HR | t(39)=1.22 |  | 0.23 | 0.193 |  |
| rMSSD | W=415 | 0.35 | 0.74 | 0.064 |  |
| LF/HF ratio | W=471 | 0.53 | 0.6 | 0.094 |  |
| Study 2 |  |  |  |  |  |
| | | df | F | p | $\eta_p^2$ |
| HR | Condition | (1.62) | 2.103 | 0.152 | 0.033 |
|  | Condition*Sex | (1.62) | 0.098 | 0.755 | 0.002 |
| rMSSD | Condition | (1.62) | 1.304 | 0.258 | 0.021 |
|  | Condition*Sex | (1.62) | 0.015 | 0.902 | 0.0003 |
| LF/HF ratio | Condition | (1.62) | 0.362 | 0.57 | 0.005 |
|  | Condition*Sex | (1.62) | 0.03 | 0.862 | 0.0005 |

Note: HR= Heart Rate; rMSSD= Root Mean Square of Successive Differences; LF/HF = Ratio of Low-Frequency to High-Frequency power.

#### EDA

In Study 1, no significant differences were observed in the tonic EDA component during the 10 minutes following HFS (t(35)=0.926, *p*= .36, Cohen’s d=0.154, BF_10_=0.27).

Anova results in study 2 revealed a significant main effect of condition on the tonic EDA component (F(1.43)=5.033, *p*=.030, n ^2^=0.105, BF_10_=2.04) and no significant interaction between condition and participants’ sex (F(1.43)=0.142, *p*=.7, n ^2^=0.003, BF_10_=0.8). Post Hoc comparisons indicated a significant difference in tonic EDA during the 10 minutes following HFS between receiving stroking from the romantic partner and being alone (*p*= .03, Cohen’s d=0.377). Descriptive results for study 2 are presented in Table 6.

**Table 6.** Descriptive tonic EDA results for study 2.

| | Sex | N | Mean ($\mu S$) | SD |
| --- | --- | --- | --- | --- |
| Dyadic | Male | 23 | 27.2 | 32.6 |
|  | Female | 22 | 30.2 | 33.0 |
| Individual | Male | 23 | 14.9 | 13.8 |
|  | Female | 22 | 21.5 | 28.7 |

Note: $\mu S$ =Microsiemens; N=Number of participants; SD= Standard Deviation

## Discussion

In this work, we explored whether different forms of CT-stimulation and social context could modulate the development of secondary hyperalgesia, an increased sensitivity to mechanical stimulation in an uninjured skin area surrounding a primary cutaneous tissue injury, that occurs due to heightened responsiveness of nociceptor neurons [4,54]. Across two studies—one comparing CT-optimal brushing to vibration, and another contrasting affective touch from a romantic partner with the absence of social support—we found no evidence that CT-activation, nor interpersonal affective touch, alters mechanical sensitization. Contrary to our main hypothesis, results from both studies show that activation of CT-receptors, both in a neutral and in a more naturalistic approach, doesn’t modulate the establishment of sensitivity to mechanical stimulation. This absence of effect was observed across different measures, such as the extent of the sensitized area, behavioral and electrophysiological responses. This suggests that neither CT-activation nor interpersonal social-affective touch is sufficient to interfere with the development of central sensitization under these conditions.

The pleasant component of touch, also known as affective touch, is mediated by CT-afferents and plays a crucial role in social interactions, being fundamental in establishing and maintaining social bonds [34,45,46]. These fibers project to brain regions that are involved in social cognition, social perception and the encoding of subjective pleasure [19,28,40]. Its analgesic potential has been demonstrated primarily in acute pain paradigms, while only a few studies have employed paradigms involving mechanisms implicated in chronic pain conditions, such as secondary hyperalgesia (SH) [16,52]. Contrary to these previous findings, our results across the two studies indicated that neither robotic nor interpersonal affective touch was sufficient to alter the development of SH. SH occurs as a result of heightened responsiveness of nociceptor neurons in the central nervous system, also known as central sensitization [4,54]. The spread of sensitization to undamaged surrounding areas is due to heterosynaptic facilitation, a form of activity-dependent synapse facilitation in which the activity of a set of synapses enhances the potentiation of other groups of adjacent synapses, previously nonactivated [31]. As a consequence, inputs from low-threshold mechanoreceptors such as Aβ-fibers, normally associated with light touch, begin to activate pain-transmission circuits [3]. All of these alterations result in heightened sensitivity to mechanical nociceptive stimuli, such as pinprick stimulation [36,48]. Our null findings did not support the involvement of the top-down or peripheral inhibitory mechanisms recruited by CT-affective touch in the spinal mechanisms of heterosynaptic facilitation that drive this heightened sensitivity to mechanical stimuli

Results from both of our studies show no effects of targeted CT-touch on any measure of sensitivity to mechanical nociceptive stimuli, including pain ratings, area of SH, or any of the psychophysiological indexes, except EDA results in the second study. This contrasts with previous work suggesting that social support (e.g., handholding) can attenuate some measurements of SH development, such as the width of the sensitized area [21]. Several methodological differences may explain this discrepancy. For instance, Jaltare KP et al. (2023) included only women and focused on social support with unrestricted partner interaction, whereas our second study included both sexes, limited the interaction to isolate touch effects, and used HFS instead of middle-frequency stimulation to induce nociceptive sensitivity.

Regarding pain levels during the HFS, we found that stroking provided by a romantic partner decreased acute pain perception, an effect not observed in the first study, where the CT-stimulation was applied “neutrally” by a robot. This difference may be explained by the emotional safety and social support inherent in partner-delivered touch, which can decrease the salience of impending noxious stimuli and facilitate analgesia [10,18,44]. Indeed, stroking prior to a noxious stimulus seems to reduce pain, whether applied by a romantic partner [42] or by a stranger [25]. Moreover, even when applied concurrently with nociceptive stimuli, analgesic effects have been observed both in acute pain paradigms and in mechanisms implicated in chronic pain conditions [16,33,52]. Although previous studies suggest that even neutral touch (e.g. robotic stroking) can attenuate pain through activation of CT-afferents or attentional mechanisms [52], our Study 1 showed that robot delivered CT-stimulation was not able to modulate acute pain. This discrepancy may be attributable to the high intensity of HFS-induced pain, which might override the more subtle modulatory effects of neutral touch. Overall, our results suggest that, while affective touch may reduce acute pain perception—particularly when delivered by a romantic partner— it does not modulate the development of central sensitization, as reflected in SH. This is consistent with findings showing that top-down modulation through attention might not be sufficient to interfere with sensitization mechanisms underlying SH [13]. The discrepancy between acute pain and SH results likely reflects the distinct levels at which these processes occur: while SH is primarily driven by spinal-level plasticity (e.g., heterosynaptic facilitation), the conscious perception of acute pain occurs at later stages of processing, more susceptible to attentional, emotional, and social contexts, like someone comforting. Thus, although affective touch has shown analgesic effects in both acute pain and some chronic pain models, it may not be sufficient to interrupt the spinal-level plasticity underlying the establishment of SH, at least under the controlled experimental conditions tested in our studies.

Across both studies, we found no differences in ECG measures. These cardiac indexes are affected by the dual sympathetic–parasympathetic innervation of the heart. Since cardiac activity reflects the net balance of these opposing branches, it often shows limited sensitivity to subtle contextual manipulations. Accordingly, measures primarily indexing sympathetic-driven arousal (e.g., HR) may respond differently from those capturing regulatory processes over time (e.g., HRV). Interpreting these patterns therefore requires considering each cardiac measure separately and with respect to the temporal phase of the paradigm to which is most sensitive. Arousal from pain or social context may increase sympathetic drive, whereas parasympathetic modulation, such as, emotional safety from partner touch, may counterbalance it and offset these effects in cardiac measures [5,27].

In contrast to the heart, the skin is exclusively innervated by sympathetic fibers [8]. In Study 2, EDA results revealed a significant increase in tonic activity when participants were stroked by their romantic partner, compared to being alone. This aligns with previous findings showing higher tonic EDA in response to touch, relative to non-CT-targeted tapping [15]. Additionally, experiences of love evoked through personalized recall have been associated with a nonspecific increase in tonic EDA [27]. The observed higher tonic EDA may reflect increased social salience in the presence of a partner, being consistent with the literature showing that tonic EDA can capture arousal in socially salient contexts [8,14,15].

The majority of studies investigating the analgesic effects of affective touch [42] or other forms of social support- such as holding hands [10,17,18,35,38] or partner presence [26]- have focused exclusively on women as pain receivers. Studies that include both sexes often fail to explore sex differences [61]. Only a previous study investigated the analgesic effects of social and interpersonal touch, comparing responses from males and females [49]. These authors found that social support decreased pain-related phasic EDA responses in both sexes, highlighting the need for further research on sex differences using additional electrophysiological measures. In our second study, we included sex as a between-subject factor, however, no significant effects were observed in any of the self-reported or electrophysiological measures analyzed. This result suggests that affective touch during the induction of central sensitization, as opposed to acute pain, may be less influenced by sex-related variables. Future studies with larger samples may help clarify whether and when sex plays a role in the social modulation of pain.

Several limitations of the current studies should be considered when interpreting the results. First, the intensity of the pinprick stimulation used (128 mN) may have limited the sensitivity of our ERP measures. Previous research has shown that cortical responses to nociceptive stimuli can reach saturation at higher intensities, reducing their ability to reflect more subtle modulatory effects [9]. While our paradigm was sensitive enough to detect changes before and after the induction of secondary hypersensitivity, the HFS may have minimized the detectability of more nuanced effects related to affective or social touch. Additionally, the variability inherent in manually applied pinprick stimulation could have introduced noise into the measurements; future studies may benefit from using robotic or automated devices to ensure consistency and precision [59]. Furthermore, our design did not control for the possible influence of the experimenter’s presence during the pinprick moments, which could have impacted participants’ pain perception [2,11,23].

In conclusion, our findings indicate that that affective touch—whether delivered in a controlled or emotionally salient social context—is insufficient to interfere with the establishment of secondary hyperalgesia. This conclusion is supported by converging evidence from two complementary studies: Study 1, with a more neutral CT-afferent activation and Study 2, with a larger sample and a more ecologically valid interpersonal context, likely associated with higher emotional and social salience. Despite these methodological strengths and the increased potential for top-down modulation in the second study, no effects were observed on any measure of mechanical sensitization.

## Acknowledgments

This study was conducted at the Psychology Research Centre (CIPsi), School of Psychology, University of Minho, supported by the Foundation of Science and Technology (FCT; UID/01662: Centro de Investigação em Psicologia) through national founds. The authors have no conflicts of interest to declare. Márcia da-Silva was supported by a doctoral research grant from the Foundation for Science and Technology (FCT) through the Portuguese State Budget (Ref.:UI/ BD/154477/2022). Alberto J. González-Villar was supported by grant RYC2023-044827-I, funded by MICIU/AEI/10.13039/501100011033 and by the FSE+. This work was supported by the EFIC-Grünenthal Grant (E-G-G) 2022.

## Supplementary Material

### N-P complex with 0.3-30Hz band pass filter

In study 1, no differences in the N-P complex (calculated as the subtraction between pre- and post-induction of SH) were observed between the robot stroking and vibration conditions (t(39)=-1.2, *p=.*24, BF_10_=0.33). ANOVA results from study 2 revealed no significant differences in the N-P complex amplitude (also calculated as the subtraction between pre- and post-induction of SH) between individual and dyadic settings (F(1,59)=0.23, *p*=.6, BF_10_=0.21). However, a significant interaction between condition and sex was observed (F(1,59)=5.5, *p*=.022, BF_10_=0.11). Post hoc tests did not reveal any significant differences between sexes within either condition. Additionally, results demonstrated weak evidence in favor of the null hypothesis for sex differences (BF_10_=0.528) (See fig.1 S.M.).

**Fig.1 S.M.**
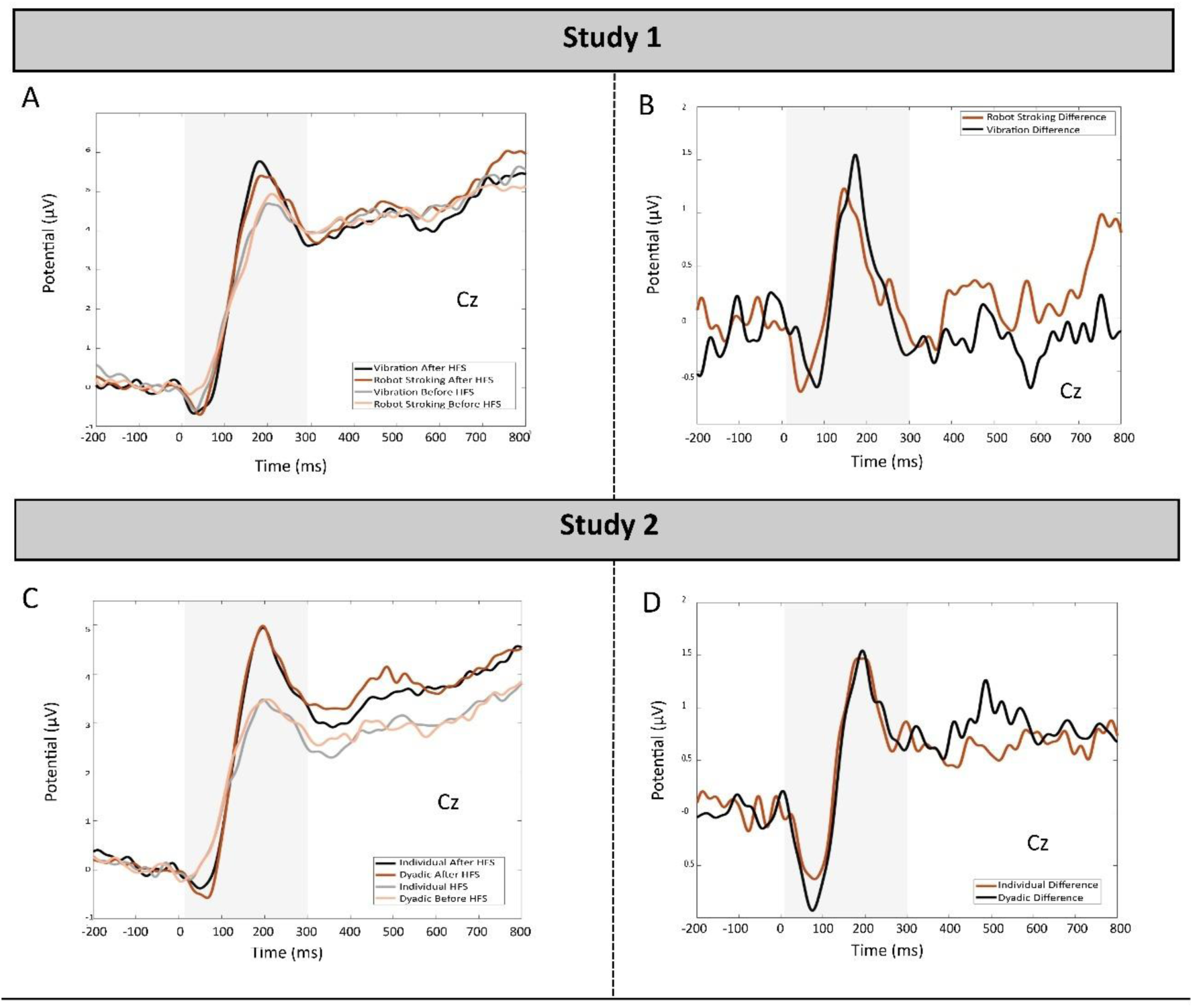
Pinprick-evoked potentials recorded over the CZ electrode in both studies with a band pass filter of 0.3-30Hz. **Study 1** (A, B): A-Pinprick-evoked potentials at baseline (T0) and post-induction (T1) for Robot stroking and Vibration conditions; B-Difference waveform (T1 minus T0). **Study 2** (C, D): C-Pinprick evoked potentials at T0 and T1 for Dyadic and Individual conditions; B- Difference waveform. HFS- High Frequency Stimulation.

## Notes

### Competing Interest Statement

The authors have declared no competing interest.

